# What do navigating animals see? Reconstructing the visual histories of desert ants

**DOI:** 10.64898/2026.10.22.760929

**Authors:** Blayze Millward, Sebastian James, Emily Baird, Michael Mangan

**Author notes:** For correspondence (FMS); (FS). These authors contributed equally to this work (Blayze Millward, Sebastian James, Michael Mangan). Department of Computer Science, The University of Sheffield, United Kingdom (Emily Baird, Michael Mangan). Department of Zoology, Stockholm University, Sweden.

## Abstract

To understand how brains translate sensory input into behavioural outputs requires a fuller account of the sensory experience of individual animals in their natural settings. Previously we developed a visual rendering software capable of reconstructing the information that insect’s compound eyes perceive (***Millward et al., 2022***). In this work, we extend this software pipeline to integrate data from animal tracking, habitat mapping, and compound eye modelling methods. The result is a new type of dataset that documents the natural foraging behaviour of desert ants (a model species for the study of animal navigation) within their natural habitat, combined with simulation tools to accurately reconstruct the visual experience of each forager as they learned to navigate their surroundings. This addresses a long-held need to create realistic training input for brain models complimented with verification tests in the same real ecologically relevant scenarios.

## Introduction

Intricate wiring diagrams of animal brains are now publicly available, produced by painstakingly reconstructing the neural connections found within the brains of various species. Complete connectomes: worms (***White et al., 1986; Veraszto et al., 2025***); fruit flies (***Winding et al., 2023; Dorkenwald et al., 2024; Takemura et al., 2025***); Partial connectomes: various insects (***Heinze et al., 2021***); mice (***The MICrONS Consortium et al., 2025***); humans (***Shapson-Coe et al., 2024***). In tandem, researchers have sought to identify the role of specific brain regions by observing which areas activate when the subjects perform a specific task (for a review see ***Jazayeri and Afraz(2017***)). An example of the success of this approach is the discovery of head-direction circuits (cells that continually track the animal’s heading direction) in mammals (***Taube et al., 1990***), birds (***Ben-Yishayetal., 2021***), fish (***Vinepinsky et al., 2020***), and invertebrates (***Seelig and Jayaraman, 2015***). By combining these data, computational neuroscientists are then able to build biologically constrained neural models to investigate what computations these brains regions perform, and the mechanisms by which they do so. Indeed, models of the insect head-direction circuit now include proposed mechanisms for altering the heading direction as the animal turns (***Cope et al., 2017; Turner-Evans et al., 2017***), integrating cues from multiple sources (***Sun et al., 2020***), and steering the animal towards a desired goal (***Stone et al., 2017; Lyu et al., 2022***).

Yet, models of deep-brain circuits (e.g. hippocampus or basal ganglia in mammals; the central complex or mushroom bodies in invertebrates) are mostly verified using highly abstracted or idealised sensory input (for a discussion see ***James et al. (2018***)). This overlooks the vital role of each species’ unique sensory organs and associated processing pathways in driving behaviour. The importance of these perception generating structures cannot be overstated with approximately 50% of brain cells of monkeys (***Felleman and Van Essen, 1991***) and insects (***Raji and Potter, 2021***) dedicated to processing visual information alone. Thus, the next frontier for neuroethology research (the discipline concerned with revealing the neural mechanisms underlying behaviour (***Ewert, 1980; Beetz, 2024***)), is to extend brain modelling studies to include more realistic sensory input.

To achieve this aim requires a new type of dataset and associated tools. This is because reconstructing the flow of information experienced by an animal requires concurrent knowledge of: the animal’s pose at a given moment in time; the sensory information provided by the environment to the animal at that location; and the information extracting properties of the animal’s sensory organs. Where possible this data should be captured in the animal’s natural habitat as they perform evolutionarily tuned tasks such as foraging for food (***Ulanovsky, 2025***).

Desert ants provide an ideal species to pioneer this approach. They are expert visual navigators, learning visually-guided routes through complex environments (***Kohler and Wehner, 2005; Mangan and Webb, 2012***) (a behaviour also observed in birds (***Biro et al., 2004***), fish (***Givon et al., 2022***), and mammals including humans (***Tamara et al., 2010; Ekstrom and Isham, 2017***)) making them ideal for comparative cognition studies. Importantly, the entire foraging paths of individual ants can be recorded in detail within their natural habitat including after experimental manipulations (e.g. ***Wehner et al. (2004***); ***Narendra et al. (2013***); ***Haalck et al. (2023***)). As these behaviours take place within a few hundred metres of the ant nest, the 3D structure of the full habitat can be mapped in detail (e.g. ***Stürzl et al. (2015***); ***Murray and Zeil(2017***); ***Risse et al. (2018***)). At the same time, the information-extraction properties of insect eyes are being revealed in evermore detail (e.g. ***Stuerzl et al. (2010***); ***Baird and Taylor(2017***); ***Juusola and Song(2017***); ***Bagheri et al. (2020***); ***Currea et al. (2023***)). And, rendering tools have recently been developed to accurately reconstruct the visual information provided by eyes to brains (***Millward et al., 2022***), which differs markedly from the output of common cameras that mimic human visual perception.

To this end, we presenta generalisable framework for next-generation neuroethology research that fuses video-based behavioural recordings with laser-based environment maps to embed high spatio-temporal behavioural data in a geometrically accurate model of an animal’s natural habitat. We then demonstrate how state-of-the-art insect eye mapping techniques and computer graphics tools can be used to reconstruct in high fidelity the visual experience of ants while they foraged. The outcome is a system by which brain models can be trained and tested with input closely aligned to that experienced by freely foraging desert ants. This provides a means to examine the fundamental principles of animal sensing, neural processing and action, and to verify and refine outcomes by comparison with real animal behaviour in the same setting.

## Materials and Methods

### Documenting the foraging lives of individual ants

The first step in reconstructing the visual experiences of individual ants is to document their movement patterns in their natural environment. Such data was presented by ***Haalck et al. (2023***) who tracked individual desert ants (species: *Catalyphis velox)* from their first forage until they found a regular feeding site in their semi-scrub habitat in southern Spain. Naïve ants were marked with specks of coloured enamel paint allowing the same ant to be tracked over multiple journeys across multiple days. The foraging journeys of marked ants were recorded using a downward-facing camcorder (1080p at 50 Hz) as they explored their surroundings freely. That is, foragers were free to move in the direction of their choice, interact with items in their surroundings, and return to the nest when they wished. However, once foragers reached approximately 8 m from the nest they were provided with a small pile of cookie crumbs representing a desirable and plentiful food source to which they returned repeatedly.

***Haalck et al. (2023***) developed custom software to convert the video recordings into trajectories describing each ant path registered to a large 2D image mosaic of the habitat surface. This dataset is unique in capturing the entire foraging lives of desert ants from their first forage, as they searched their environment, encountered food, brought it home, returned to the feeder site, and were subjected to experimental manipulations (captured at the nest and released at the feeder to assess visual memory). These high spatio-temporal resolution path data are ideal as a basis for reconstructing the sensory stimuli experienced by an animal completing an ecologically important task in the wild. Throughout this work we focus on the foraging paths of 4 ants from the original dataset (Ants 3, 6,11,12) which are representative of the general foraging behaviour of the colony, and provide ample data to demonstrate the potential for our approach (***Table 1,*** ***Figure 1****A*).

**Table 1.** Ant Trajectory Data.

| Ant | Colour Code | Data |  |  |  |  |  |  |
| --- | --- | --- | --- | --- | --- | --- | --- | --- |
| 3 | Blue | $R1_1$ | $R2_2$ | $R3_3$ | $R5_3$ | $ZVF$ | | |
| 6 | Green | $R1$ | $R2$ | $R3_1$ | $R4_2$ | $R5_2$ | $ZVF$ | |
| 11 | Pink | $R1$ | $R2$ | $R3_1$ | $R4_2$ | $R5_3$ | $R6_3$ | $ZVF$ |
| 12 | Orange | $R1_1$ | $R2_1$ | $ZVF$ | | | | |

Rows describe trajectory data from the original dataset (Haalck et al., 2023) that we use in this study. Ants retain their numbering as in the original study for consistency. Each ant is given a colour code that is maintained in all trajectory plots throughout this paper. Entries with the prefix $R$ indicate a foraging “run” with the number indicating position in the sequence (e.g. an ant’s first forage is $R1$ , second forage is $R2$ ). Runs where ants reached 8m and were rewarded with cookie are given a subscript with each entry indicating a feeding location (some ants didn’t return to precisely the same location). $ZVF$ trajectories are from the test condition whereby ants were captured just before entering the nest (hence zero-vector, or $ZV$ ) and released at the just visited feeder location ( $ZVF$ ). In plots, all $ZVF$ data is plotted in white to distinguish them from normal foraging. There is no entry for Ant3R4 as in Haalck et al. \(2023\).
Source Data: Dataset: Reconstructing the visual histories of desert ants (Mangan et al., 2026).

**Figure 1.**
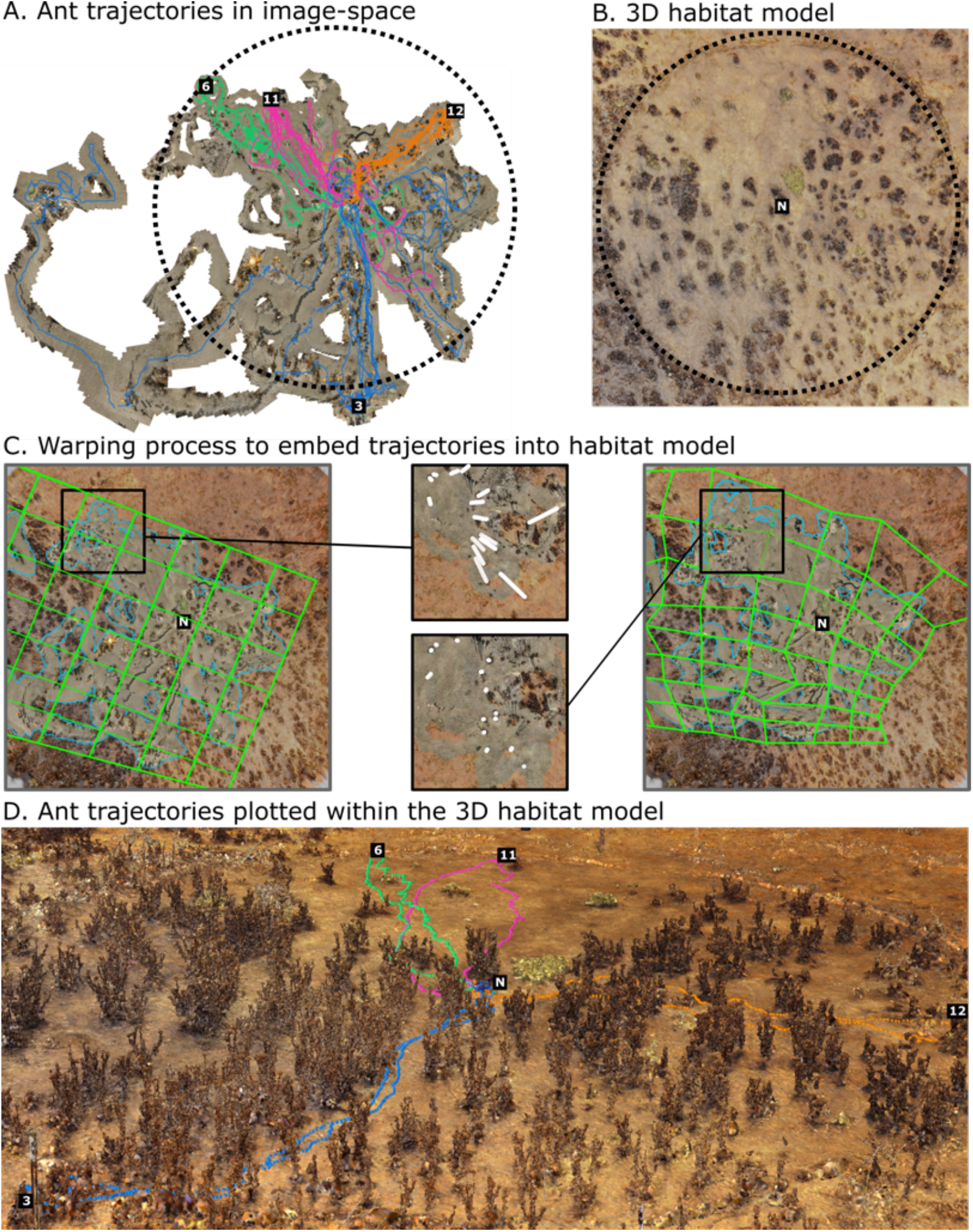
Translating image-based trajectories into the 3D habitat model. (A) The complete foraging paths of 4 desert ants are shown (Ant3: blue; Ant6: green; Ant11: pink, Ant12: orange) overlaid onto the image-space model as in ***Haalcket al. (2023****). N* shows the nest location, numbered squares show each ant’s final feeding site, and the dotted circle shows the experimental recording area with an 8 m radius. (B) An overhead projection of the habitat model created from laser scans (***Risse et al., 2018***). (C) Overview of the warping process used to translate trajectories from image-space to habitat-model space. The leftmost image shows: the pre-warped warpmesh as a green grid; the image mosaic (blue outline) (***Haalck et al., 2023***) overlaid into an overhead image taken from the habitat model (background) ***Risse et al. (2018***). The central images show a patch of the scene pre (upper insert) and post (lower insert) warping. The white lines show the distance between the same manually identified locations in the image and habitat models. The rightmost image shows the aligned image model (outlined in blue) created as a function of the adapted warp-mesh (overlaid in green). (D) The final foraging path of each ant before displacement tests is plotted within the habitat model (Ant3R5: blue; Ant6R5: green; Anti 1R6: pink, Anti 2R2: orange). *Source Code & Data:* [A-C Image Alignment](https://github.com/BrainBeatsBrawn/image-aligner); [D: Plot trajectory data](https://github.com/BrainBeatsBrawn/antpov).

### Mapping the ant habitat

Desert ants are primarily visual navigators, utilising cues such as the silhouette of the surrounding skyline for guidance (***Fukushi, 2001; Graham and Cheng, 2009***). Thus, we require a map of the spatial configuration of objects in the animal’s habitat (ground, bushes, rocks) if we are to reconstruct the visual information incident to each forager’s eye. We make use of a 3D model of the ant habitat created from 56 laser scans captured across the foraging site from which the above behavioural data was collected (***Risse et al., 2018***). The resultant mesh model provides an accurate representation of the colour and 3-dimensional shape of the ground surface and the objects in the foraging area, allowing photorealistic visualisation of the environment (***Figure 1***B, D). Given that the habitat underwent little structural change over the recording period, this model can be used for reconstructing the visual experience of all ants from the dataset.

### Embedding ant trajectories into the habitat model

As the ant trajectories and 3D habitat model were generated using different sensors (camera vs laser scanner) with distinct processing pipelines, it is important to ensure that they are fully aligned before any visual reconstructions are attempted. That is, confirming that the location of an ant in one corresponds to the same location in the other. We take the 3D habitat model generated by the laser scans to be metrically accurate given that depth errors fall below 2.2 mm (1 mm systematic error; 1.2 mm precision error) for measurements taken within 10m of the scanner (Z+F IMAGER 5006i in ‘high resolution’ mode) (***Zoller + Fröhlich GmbH, 2012***). Moreover, the 56 independent scans were registered using several external markers ensuring excellent alignment. High accuracy is supported by ***Stürzl et al. (2015***) who mapped insect habitats in Australia using the same hardware and reported high visual similarity between simulated and real panoramic images generated from the same location.

In contrast, we expected to see some drift in the absolute metric positions of the ant trajectories which were captured from an unstabilised, downwards facing camcorder. Such data creates multiple challenges for accurately tracking the camera pose (a key step in producing trajectories): visually sparse parts of the environment (e.g. areas of sand) don’t provide easily trackable visual features; re-projection errors increase when the camera tilts; lack of depth information makes it hard to estimate camera velocity or accurately model the 3D structure of the ground; and the abstraction of the 3D ground surface to a 2D background image is an approximation (***Haalck, 2023***).

We thus created a pipeline to transfer all trajectories from tracking-space (generated from video) into the habitat model-space (generated from laser scans). We formulated the problem as aligning the tracking-space image mosaic onto which all ant trajectories are registered (***Figure 1***A) onto an orthographic projection generated from the 3D habitat model (***Figure 1***B). We created a GUI-driven program that overlaid a semi-transparent version of the tracking-space image over the habitat-space image from which misalignments in the location’s vegetation can be identified (***Figure 1****C* - left). At a macro-level, image and habitat models are generally consistent, but inconsistencies arise when looking at the precise position of objects (***Figure 1****C* - centre). Hand labelling the locations of the same visual features (correspondence pairs) in both images found an average displacement of 59.13 cm (std: 29.16 cm, n=97), with the magnitude and orientation of misalignments varying across the environment. We thus implemented a multi-patch-warping technique (inspired by the SplitWarp algorithms in Adobe Photoshop (***Lacroute and Levoy, 1994***)) whereby different subregions of the environment could be manually warped to reduce the distance between correspondence pairs. The multi-patch warping method allows for locally accurate alignment while avoiding discontinuities at the boundaries between subregions (akin to stretching an elastic sheet over pegs). By manually warping 64 local subregions in turn, we reduced correspondence distances to 11.04cm (std: 8.36cm) (***Figure 1****C* - right). We note that adding additional subregions didn’t improve results as manually identifying correspondence pairs becomes increasingly difficult as the subregions decrease in size. Additional improvement may also require the ability to merge parts of the image mosaic which is not possible using the current method. Regardless, the current data remains sufficient for the needs of this study, and future improvements to the transformation matrices could easily be incorporated.

The resultant warpmesh was used to transform the 2D trajectory points from image-model-space (***x***^*i*^, ***y***^***i***^) into their corresponding 2D co-ordinates in habitat-model-space (***x***^***h***^, ***y***^***h***^). We estimated the 6D pose (***x***^***h***^, ***y***^***h***^, ***z***^***h***^, ***ϕ***^***h***^, ***θ***^***h***^, ***ψ***^***h***^) for each datapoint by defining the ***z*** coordinate to be 2 mm above the habitat-model’s ground surface *(C. velox* foragers stand 1-2 mm tall (***Cerdá, 2001***)), and derive roll (***ϕ***), pitch (***θ***), and yaw (ψ) angles from the velocity vectors linking 3D trajectory points, together with the gradient of the ground surface. We note that the resultant trajectories include occasional gaps (e.g. while an ant was invisible while climbing in a bush), or small positional jumps in the camera tracking. Notwithstanding, the presented data documents in unprecedented detail the entire foraging history of individual ants, now embedded within a highly accurate model of their habitat, providing the necessary information to recreate their visual experiences (***Figure 1***D, ***Table 1***).

### Mapping the light sensing elements of *C. velox* eyes

With the pose of foragers defined relative to sources of visual information in the environment, we now need to understand what part of the scene their eyes capture data from. Desert ant vision is dominated by a pair of apposition compound eyes formed by a 3D lattice of interlinked, independent light sensing elements known as ommatidia (***Land and Nilsson, 2012***). Thus, a first approximation of the light capturing properties of their eyes can be garnered from the central viewing vector of each ommatidial lens (known as a facet). Given the enormous variety of compound eye configurations across species (size, shape, number of ommatidia etc) we completed a study to map the location, size, and central viewing angle of all facets in a *C. velox* eye.

Specifically, a *C. velox* forager was captured from the same foraging site in which the field study described above was conducted before being preserved in alcohol and transported to the lab for analysis (credit to Dr. Sebastian Schwarz for specimen capture, fixing and shipping). The specimen’s head and right eye were then scanned using X-Ray microtomography methods as described in ***Baird and Taylor(2017***); ***Taylor et al. (2019***, 2020) (see ***Figure 2****A*) (credit to Dr Gavin Taylor for assistance in the scanning process). The position, diameter and central axis of each ommatidium was then estimated using a process adapted from ***Taylor et al. (2020***) (see ***Millward(2025***) for detailed description). In brief, the region of the scan representing the animal’s right eye was manually sectioned from the head and converted into a 3D surface (***Figure 2****A*). Each facet was then semi-manually identified and outlined on the surface (each facet is visible as a small bump) providing a means to estimate each facet’s 3D position, its diameter, and its central optical axis which we define as the normal projection from the facet surface (***Figure 2***B). The eye surface was then registered back to the head surface and mirrored to the left side to create a pair of identical compound eyes positioned appropriately on each side of the animal’s head (***Figure 2***C).

**Figure 2.**
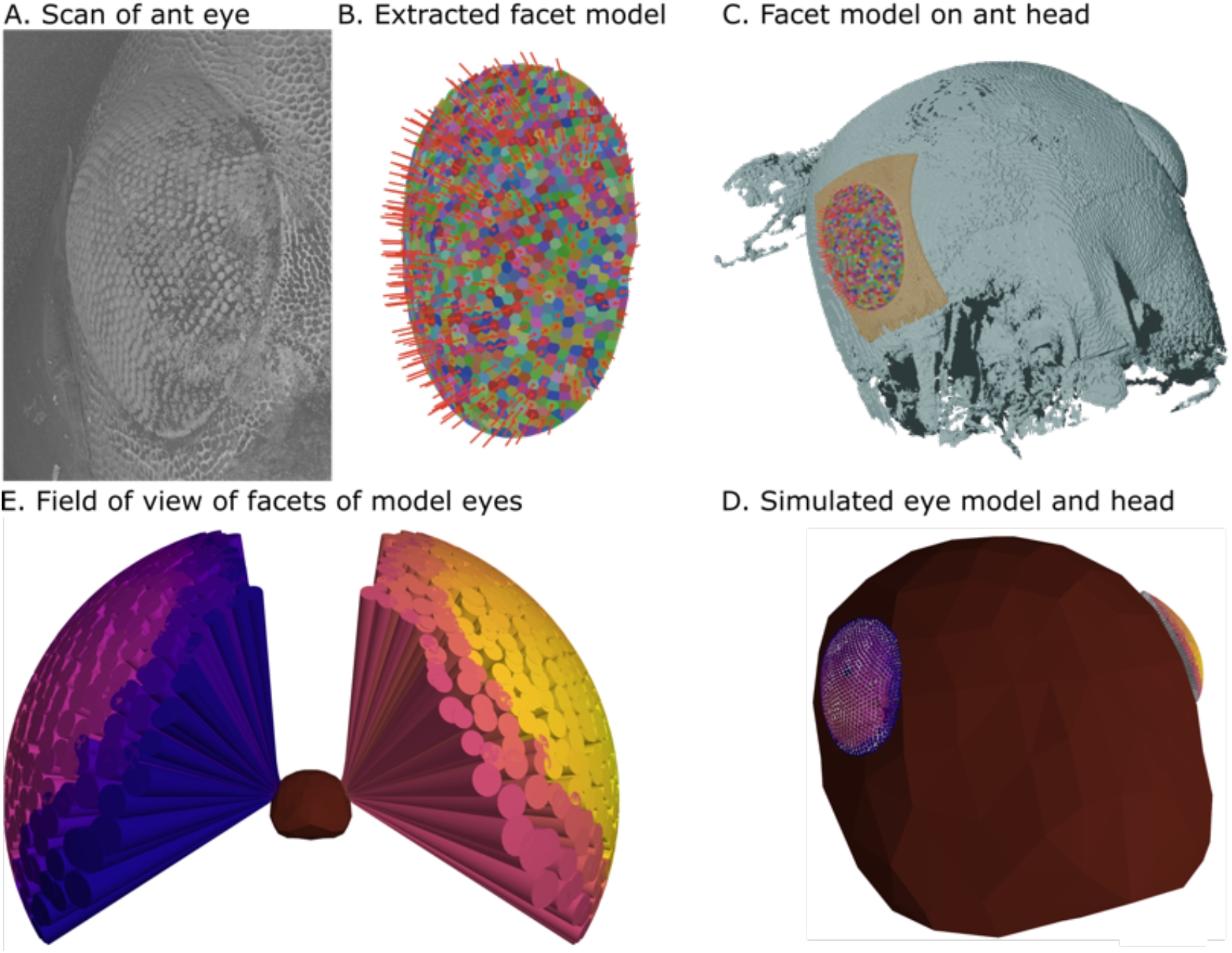
Creating biologically realistic eye models. A. X-ray microtomography scan of the right compound eye of a *C. velox* forager. B. Visualisation of the individual facets extracted from the micro-CT data. Facets are individually coloured, with red lines showing their estimated central optical axis. C. Visualisation of the facet eye model registered to the head scan. D. Visualisation of the same ant head and eyes using our visualisation tools. E. Cones projecting from each facet in the simulated ant eye model depict each facet’s estimated field of view (estimated from measurements of the facet diameters). The ant head is shown facing forward towards the viewer. *Source Code & Data:* [A-C Ant Eye Mapping and Modelling](https://github.com/BrainBeatsBrawn/scan-to-oces-pipeline); [D-E: Visualise eye data](https://github.com/BrainBeatsBrawn/antpov).

Our analysis identified 915 individual facets in the sample eye. Interommatidial angles (defined here as the average angle between the central optical axis of each facet and its neighbours) ranged from 1.4°-7.8° (mean 3.6°). This method does not allow for a direct measurement of the ommatidial acceptance angle (***Taylor et al., 2019***). These results are broadly in line with other desert ants species: *Cataglyphis bicolor (****Wehner et al., 1971; Eheim and Wehner, 1972; Menzi, 1987; Zollikofer et al., 1995***) has 600-1200 facets per eye (***Menzel and Wehner, 1970***), with interommatial angles ranging between 3.71°-6.69° (***Zollikofer et al., 1995****); Melaphorus bagoti* has 420-590 facets per eye (***Schwarz et al., 2011***) with interommatidial angles ranging between 3.7°-6.4° (mean 3.7°) (***Schwarz et al., 2011***).

### Rendering the ant’s view of the world

We now require a means to simulate the light incident to each forager’s eyes within the habitat model. To render the ant’s eye view we used CompoundRay (***Millward et al., 2022***): a software designed using modern raycasting techniques to simulate compound eye vision. The software takes a specification of a compound eye as a set of ommatidial positions, directions, acceptance angles and ‘focal point offset’ (an optical length). It is also given the habitat model as a glTF-encoded 3D scene model. The software then loads the scene (triangle vertices, vertex colours, vertex normals, a sky model and any textures) and sets up a ‘virtual camera’ for each ommatidium. We here call the set of all ommatidia the *’compound-ray camera’* (in this study, the compound-ray camera comprises both ant eyes). It then performs a ray-casting operation to determine the colour detected by each ommatidium, from any location within the habitat model (e.g. the colour of sky, earth, or bush as in the real environment). The resultant data may be stored into a file, displayed in a visualization, and supplied as input to computational models for closed loop control. We used the library *mathplot (****James et al., 2026***) to produce perspective visualizations of the environment, a third person perspective of a simulated ant in the environment, and the visual input that would be perceived by the ant’s eyes. Mathplot also manages a navigation mesh that allows the simulated ant to traverse the ground of the habitat model. Note that the visualisation of the environment and ant is independent from CompoundRay whose task is to compute the colour data incident to the eye model.

***Figure 2****D* shows the model ant eyes positioned on an ant head which was used throughout this work to reconstruct the ant’s visual input, and ***Figure 2***E shows the light sampling cone of each ommatidium giving an indication of the ant’s overall field of view.

## Results

### Reconstructing the visual experience of individual foragers

We now possess all requisite information to reconstruct the visual experience of the ant foragers in their natural habitat. ***Figure 3*** shows reconstructed views from two important occasions in the foraging life of Ant3: the first time she exited the nest to forage (***Figure 3****A*); and the first time she found cookie (***Figure 3***B). These are isolated snapshots of her foraging life, but the provided dataset and associated rendering tools presented allow all views experienced along each of her foraging journeys to be reconstructed in full. As an example, see the linked video [Ant3 view reconstruction] which shows the views that Ant3 would have experienced throughout that first forage. The video is shown with time-synced positioning data in the 3D model as well as the original camera recordings, from which it is possible to identify the same plants in both. For example between 0:32-0:34 the ant approaches a large, low-lying plant that is clearly visible in both the habitat model and the original video. Notice that as the ant disappears under the foliage, this is reflected in the simulated eye data. We also note that there remains some residual misalignment (as reported above): between 1:38-1:58 when the track should remain inside not outside of the vegetation. Otherwise, the pipeline works remarkably well. This process can be repeated for all trajectories of each ant in our dataset allowing for the reconstruction of their unique visual histories as they explored their environment, and learned the location of the plentiful food source. Indeed, to the best of our knowledge this is the first time that the visual history of any animal has been documented in a manner that allows it to be reproduced at a later date.

**Figure 3.**
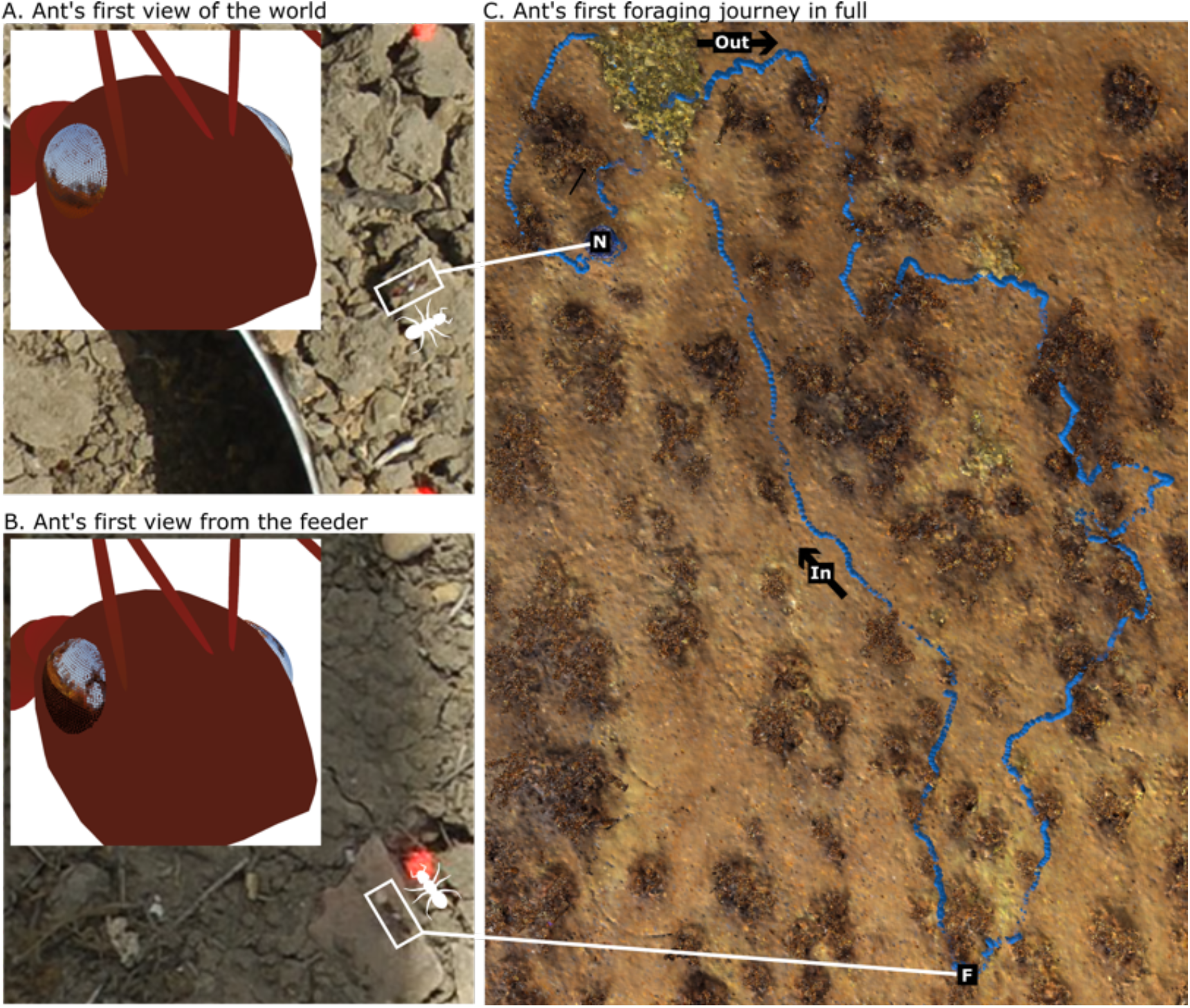
Reconstructing the views experienced by a foraging desert ant. A. The insert shows a reconstruction of the visual input that would have been experienced by Ant3 as she exited the nest area for the first time (visual input projected on the ant eye/head model). Background images show the frame from the video dataset (***Haalck et al., 2023***) as she left her nest for the first time (ant outlined by white box, orientation shown by ant icon). The corresponding position in the habitat model twin is shown by the white line. B. The same visual reconstruction but rendered from the location that Ant3 first collected cookie. C. The completAnt6R5: green; Ant11R6: pink, Ant12R2e first foraging journey of Ant3 is shown. *N* shows the nest location; the outward section of the journey is labelled *Out; F* shows the location at which she found cookie, and the subsequent journey home is labelled *In*. *Source Code & Data:* [A-C: Plot ant data and visualise visual renderings](https://github.com/BrainBeatsBrawn/antpov).

These *high fidelity ant eye reconstructions* provide a distinct view of the visual information available to the ant’s brain. Firstly, the *biologically constrained views* differ markedly from the cylindrical panoramas typically used to approximate insect vision in modelling studies (***Figure 4***A). The real eye model includes blind spots, distancing between eyes, non-uniform spatial sampling, and complex 3D structure. Eye geometry is already known to impact the form of optic flow information available to insect brains (***Misiek et al., 2026***), and shapes the underlying neural mechanisms (***Zhao et al., 2025***). By supplying this enhanced input to new models of the insect optic lobes (recently mapped in detail by ***Nern et al. (2025***) for flies) we can better hope to reveal the role and function of those circuits, and the data that they provide to deep brain areas.

**Figure 4.**
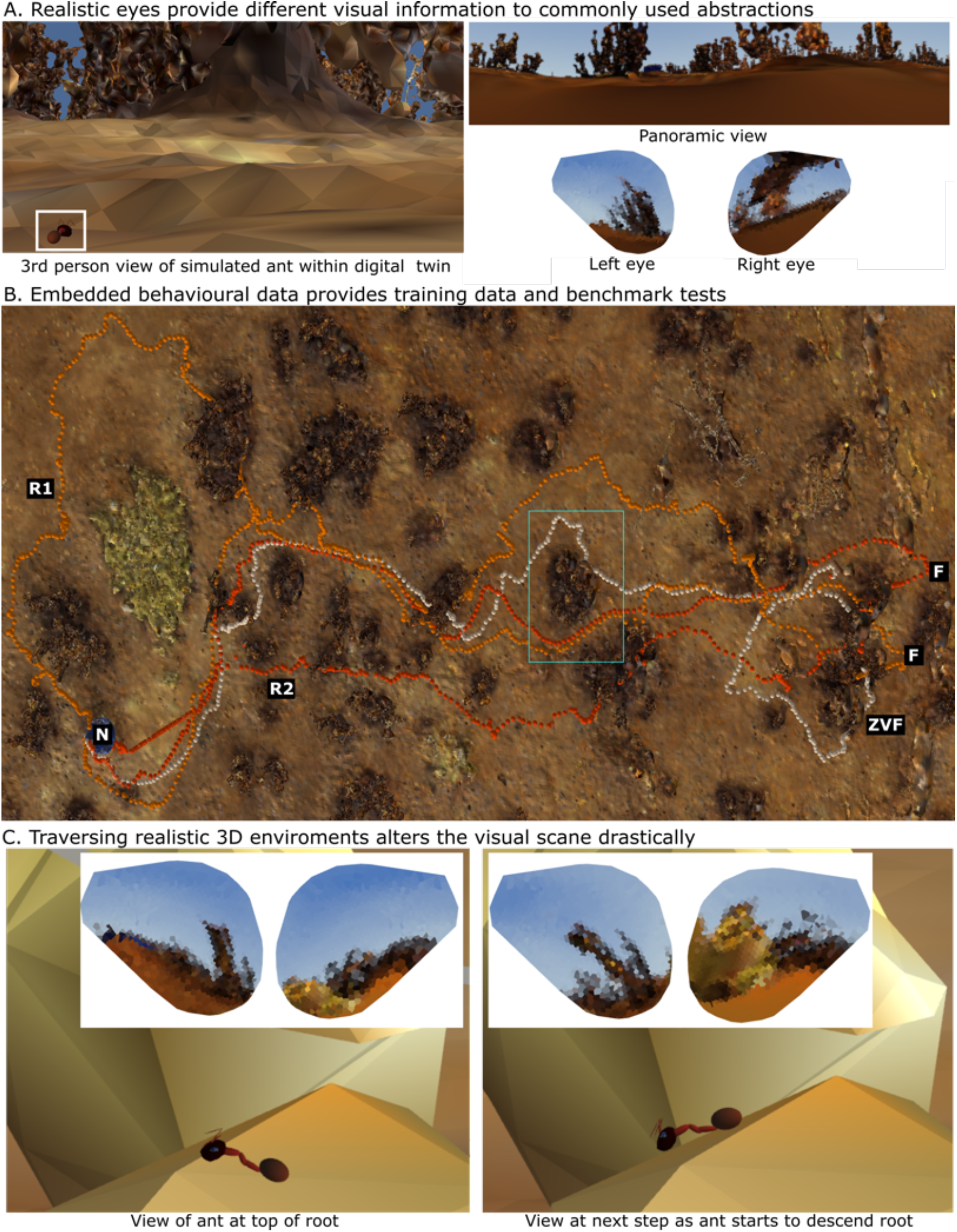
Challenges of real environments. A. Left-side insert shows a simulated ant (white box) in the 3D world. Right-side images show a panorama commonly used to simulate insect vision (360° by +/-45° around the horizon) and the output of the biologically constrained eye model from the same location. B. The first foraging paths of Ant12 are shown in orange *(R1* & *R2); N* is the nest location, and *F* shows Ant12’s feeding locations. Ant12’s path after being released at the feeder is shown in white (labelled *ZVF)*. After an initial loop, she closely followed her previous homeward paths aside from passing a bush on the opposite side (cyan box). C. A simulated ant is shown at two data-points on a real route that are very close in space and time. The uneven ground surface creates very different visual input (shown in inserts). *Source Code & Data:* [A-C: Plot ant data and visualise visual renderings](https://github.com/BrainBeatsBrawn/antpov).

This is further supported by the *high spatially-temporally accurate behavioural data* available in the dataset, which inherently structures the visual data stream. Indeed, it is possible to note differences in the visual input during the first foraging journey of Ant3 as shown in the video above. While foraging for food (e.g. 08:24-08:30) Ant3 changes heading constantly causing large and rapid lateral scene change, whereas when travelling home with cookie (e.g. 09:29-09:34) she follows a smoother path which keeps visual information aligned for longer. ***Haalck et al. (2023***) reported that homewards trajectories are learned after a single traverse, while outward trajectories are refined over time, and this difference in input data may play some role in that outcome.

The same dataset also provides well defined training and test conditions, allowing brain model performance to be evaluated against real animal data. For example, ***Figure 4****B* - ***ZV F*** demonstrates an ant successfully returning to the nest, after been displaced back to the feeder (all ants tested achieved a similar outcome). That is, in the absence of path integration assistance, they were able to recognise and retrace their familiar path home (equivalent to the kidnapped robot problem in robotics (***Engelson and McDermott, 1992***)). Due to the high-spatio-temporal resolution of the trajectory data it is possible to perform targeted analysis in instances where behaviours diverge in the animal data (e.g. ***Figure 4****B* - *cyan box* when the ***ZV F*** ant performed differently from normal homing), and between animal and model data. The dataset can also be repeatedly mined for additional insights when they arise. For example by reanalysing data for identify previously unconsidered behavioural motifs that may improve model performance. That is, the approach provides researchers with a modern, expandable, long-lasting research resource to effectively study how brains generate complex behaviour in the wild.

Finally, the *geometrically accurate 3D habitat model* offers new challenges for researchers. The habitat model’s detailed map of complex terrain over which ants must traverse (gravel, rocks, bushes) causes significant changes to the visual input experienced even over a very short period of time (e.g. ***Figure 4****C*). Previous studies have shown that state-of-the-art pixel-based visual matching algorithms are compromised when subject to pitch (***Ardin et al., 2015***), and thus mechanisms to improve tilt-tolerance will be required in future models (e.g. ***Stone et al. (2016***); ***Gkanias et al. (2019***)) if the robust behaviour of desert ants is to be replicated. This is just one example of the way that accurate 3D environmental models can capture the complexities of the real world, which can challenge and inspire new hypothesis (expanded in ***section*** Discussion). If models are to truly explain behaviour they will ultimately need to overcome the same real-world problems as the animals do, and datasets such as that presented encapsulate many of those challenges.

## Discussion

Previously, we presented an open-source rendering software *(CompoundRay* - ***Millward et al. (2022***)) capable of reconstructing the visual information available to compound eyes in abstract scenarios. Here, we integrate that software into an extended methodological pipeline that links real-world behavioural data with the ability to reconstruct the sensory stimulus of animals. Specifically, we combined data from video-based animal tracking and laser-based environment mapping tools to create a dataset that embeds real foraging data (both spontaneous and after experimental manipulation) into a photorealistic 3D model of the animal’s natural environment. We then used modern compound eye mapping techniques to create a geometrically accurate model of the ant eye to constrain the parameters for our rendering software. Together, this allowed reconstruction of the visual histories of individual desert ant foragers as they explored, learned, and responded to experimental manipulations in their complex desert habitat.

Capturing behavioural data in this way offers a number of benefits for neuroethology researchers. Firstly, the datasets generated inherently embody detailed descriptions of the experimental paradigm and preserve unprocessed behavioural data making it easier to replicate experiments and compare results across studies/species (***Korbmacher et al., 2023***) while aligning closely with FAIR data principles (***Wilkinson et al., 2016***). Compare this to the challenge of replicating studies from abstract, textual descriptions of experimental setups written into Methods sections of manuscripts (***ADMC, 2019***). We expect the accessibility of these tools to improve drastically in the near future as the cost and complexity of hardware and data-processing methods fall. Indeed, it’s already possible to create photorealistic environment models using consumer cameras and depth sensors (e.g. ***Mildenhall et al. (2021***); ***Kerbl et al. (2023***)) rather than the costly laser scanners used in ***Risse et al. (2018***). Similarly, methods to capture detailed behavioural data of animals over ever expanding ranges in the wild are advancing at pace (e.g. ***Haalck et al. (2020***); ***Smith et al. (2021***); ***Koger et al. (2023***);

### Stentiford et al. (2026); Daniel et al. (2026))

These development will only accelerate with adoption, alignment around best practices, and feedback to technologists creating a virtuous cycle where both research outcomes and methods advance together.

Secondly, by closing the loop between animal movement in the wild and reconstruction of the sensory stimuli experienced by an animal, we provide an essential bridge between brains and behaviour. Currently, brain models are prototyped in abstract simulations that rarely present realistic sensory stimulus compared to natural conditions. Moreover, it remains cumbersome to verify brain models in embodied behavioural tests that accurately replicate the challenges faced by animals (usually requiring robots and custom sensory apparatus). Our approach brings the complexity of the real-world into simulation, allowing analysis of the sensory input by researchers, provision of realistic input to brain models, and verification and refinement of model outputs through direct comparison to real animal behaviour. We note that a similar approach (oft termed Real-to-Sim-to-Real or Digital Twins) is becoming increasingly pervasive in robotics research where it has significantly reduced development time, and translation reliability from simulation to real world deployment (***Mazumder et al., 2023; Torné et al., 2024; Dong et al., 2025; Zhu et al., 2025***).

The approach presented is both extendable and generalisable providing a level of future proofing for data captured in this way (that is preserving the usefulness of data for longer). An obvious first extension would be to add more realistic lighting through sky models (e.g. ***Gkanias et al. (2019***)) presenting new challenges to brain models to overcome such as shadows, clouds, chromatic gradients and even new visual cues such as polarisation. This could coincide with advancing the biological realism of eye models by adding advanced photoreceptor models (e.g. ***Juusola and Song(2017***)), microsaccades (e.g. ***Juusola et al. (2017***); ***Mansour et al. (2026***) and more. And there is the potential to map and integrate other modalities of sensory cues such as odour (***Buehlmann et al., 2015***), magnetism (***Fleischmann et al., 2020***), and even vibrations (***Buehlmann et al., 2012***), while incorporation of physics into simulations would allow the influence of wind on navigation to be added (***Müller and Wehner, 2007; Desai et al., 2026***). While we have focussed this work has been desert ant behaviour, and compound eyes, the system is highly flexible and generalisable. For example, CompoundRay could be adapted to model the curvature and non-uniform sampling (e.g. peripheral vs foveated vision) found in mammallian eyes and extended to various experimental settings. There is also the potential to integrate with other modern neuroscientific tools such as virtual reality (see ***Kócsi et al. (2020***) for review) to capture high precision behavioural data such as oscillations (***Clement et al., 2025***), and neural responses in behaving animals (e.g. ***Seelig and Jayaraman(2015***)) when tested in environments closely resembling their natural habitat.

The potential for emerging tools such as those demonstrated to accelerate neuroethology research is enormous. However, to reap those rewards, a community effort is required to define common standards, tooling and datasets akin to that which happened as connectome mapping technologies were increasingly adopted across laboratories (e.g. ***Shinomiya et al. (2011***); ***Ito et al. (2014***)). We are now in a position to peer out at the world through the eyes of a navigating animal, and link what we see to brain models to inspire and challenge our hypotheses of how brains drive behaviour.

## Data and Code Availability

*Data:* [Dataset: Reconstructing the visual histories of desert ants](https://doi.org/10.15131/shef.data.32860832) (***Mangan et al., 2026****). Code and data usage instructions:* BrainsBeatsBrawn.github.org.

## Acknowledgments

We extend our gratitude to Dr Sebastian Schwarz for capture, fixing and shipping of forager used for eye mapping; to Dr Gavin Taylor for assistance in generating eye scans; and to Rebecca Fuidge for assistance of the manual labelling of eye facets.

